## Supplementary Information for "A Synergistic Workspace for Human Consciousness Revealed by Integrated Information Decomposition" (from another version of this preprint: version 4, 2024-02-08)

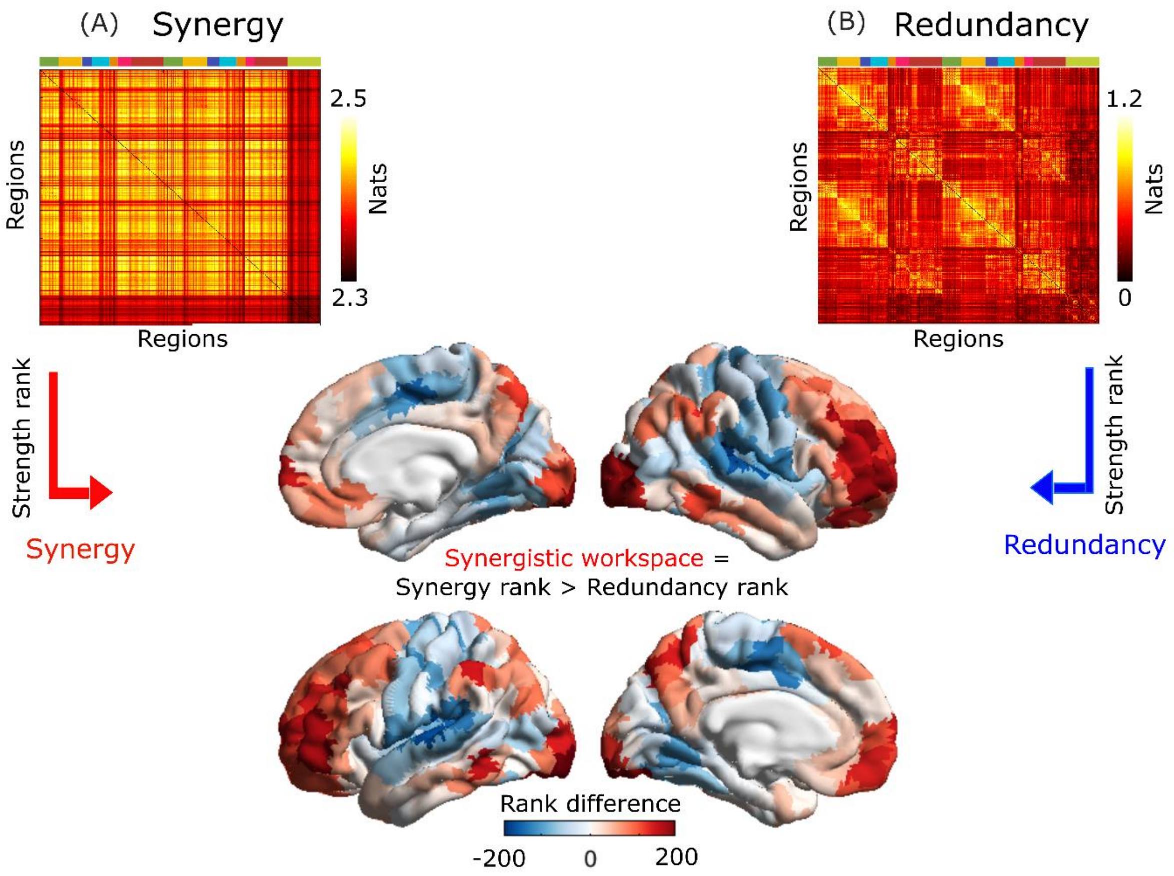


**Figure S1. Identification of the synergistic workspace.** Following Luppi *et al.,* (2022), regions of the synergistic workspace (red) are identified as those regions whose rank, in terms of strength of synergy with the rest of the brain, is greater than the corresponding strength rank for redundancy.

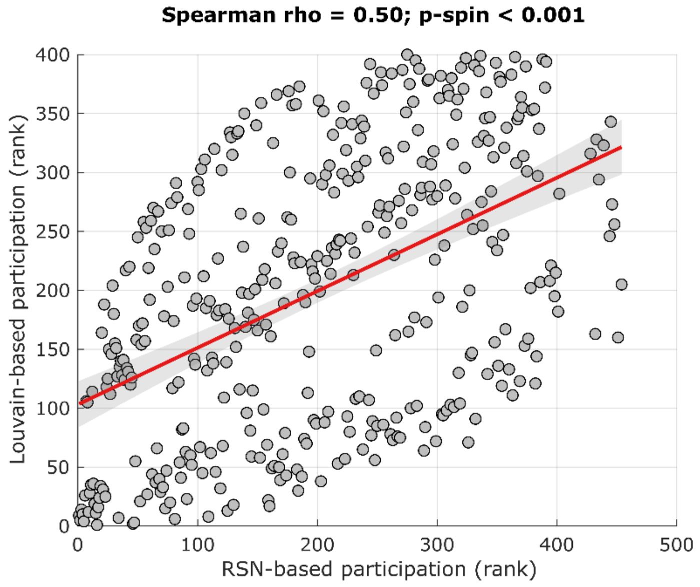


**Figure S2. Significant correlation between regional participation coefficient computed with modules defined as resting-state networks (X-axis), and with modules defined from Louvain modularity detection (Y-axis).** Significance is assessed against a null distribution of 10,000 autocorrelation-preserving maps.

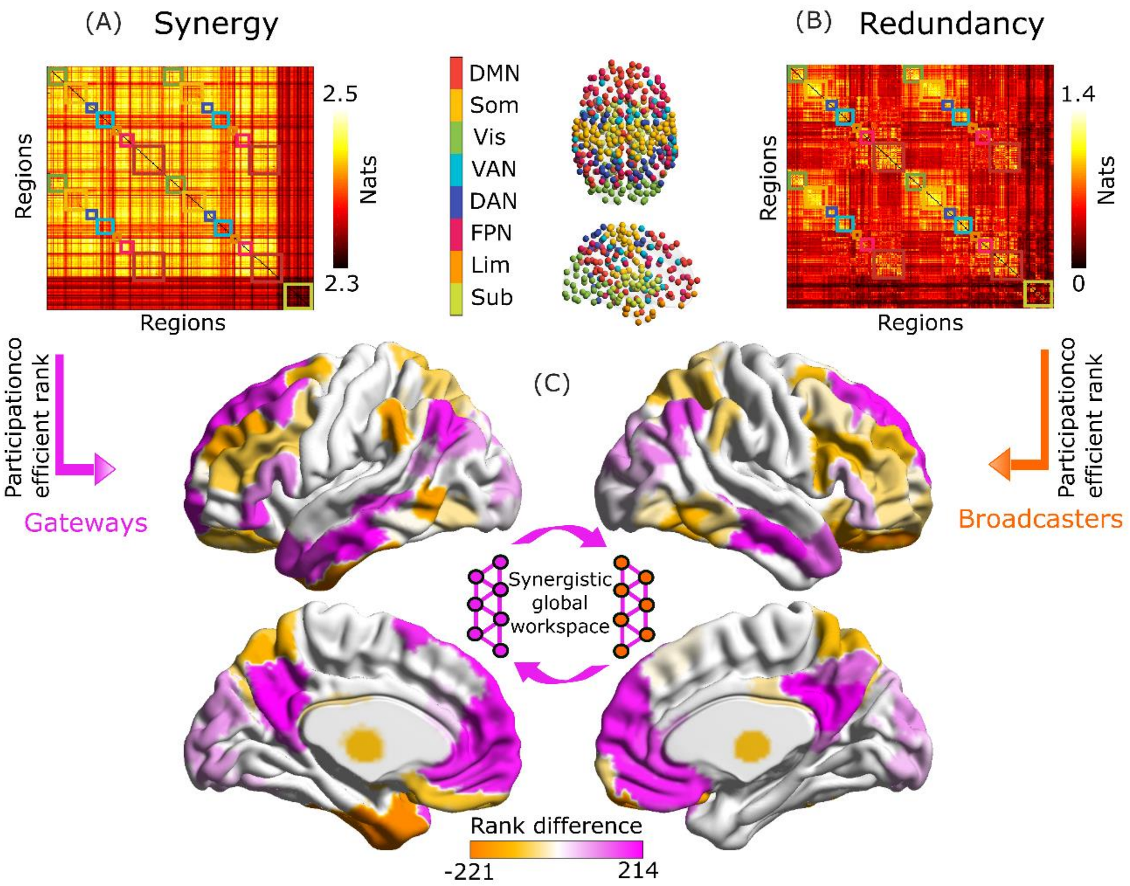


**Figure S3. Identification of workspace gateways and broadcasters is robust to node definition.** Gateways (violet) and broadcasters (orange) identified on the Schaefer-232 parcellation.

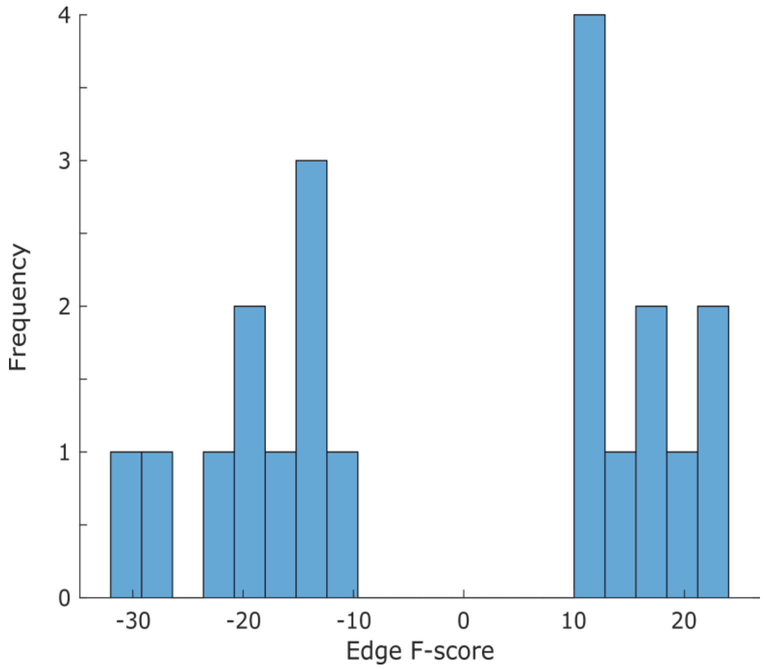


**Figure S4. Histogram of significant connectivity changes.** Distribution of F-scores for brain connections that are consistently and significantly increased (positive) or decreased (negative) during unconsciousness.

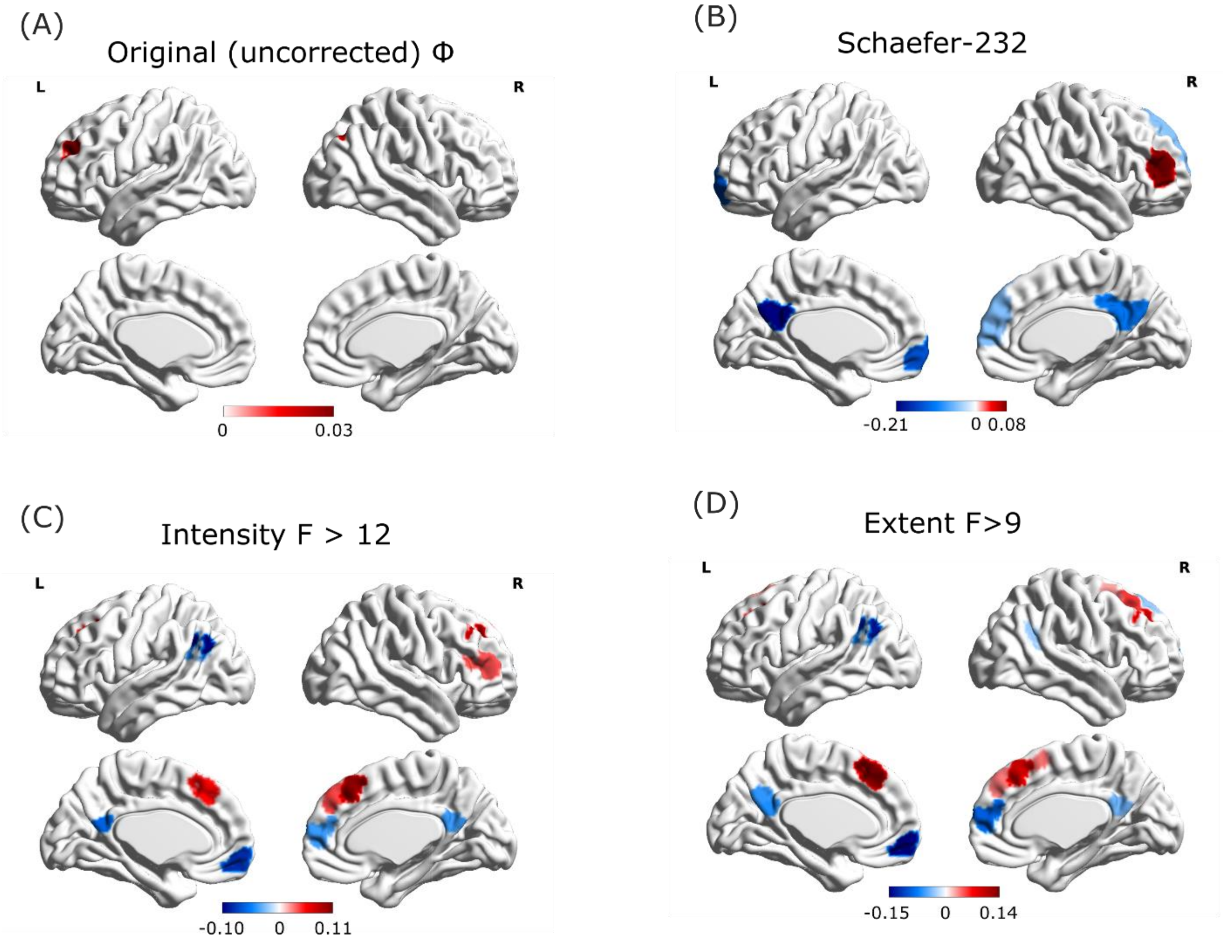


**Figure S5. Results of alternative analysis choices.** Brain regions exhibiting overall increases (red) and decreases (blue) in integrated information exchange, when comparing patients with disorders of consciousness with awake healthy volunteers, propofol anaesthesia with pre-induction wakefulness, and propofol-anaesthesia with post-anaesthetic recovery. (A) When the original formulation of whole-minus-sum Φ (Balduzzi 2008) is used, no consistent reductions are observed: only increases. (B) Results with revised Φ are robust to the use of an alternative parcellation scheme (Schaefer-200 cortical ROIs plus Tian-32 subcortical ROIs). (C) Results with revised Φ are robust to the use of a more stringent threshold value (F > 12). (D) Results with revised Φ are robust to the use of alternative threshold definition (extent vs intensity).
